# Stability Criterion for the Assembly of Hybrid Lipid-Polymer-Nucleic Acid Nanoparticles

**DOI:** 10.1101/2022.02.06.479316

**Authors:** Juan L. Paris, Ricardo Gaspar, Filipe Coelho, Pieter A. A. De Beule, Bruno F. B. Silva

## Abstract

Hybrid lipid-polymer-nucleic acid nanoparticles (LPNPs) provide unique delivery strategies for nonviral gene therapy. Since LPNPs consist of multiple components that can undergo different pairwise interactions between them, LPNPs are difficult to prepare and characterize. Here we demonstrate that the interaction between the polycation (polylysine) and DNA is robust through an innovative implementation of fluorescence cross-correlation spectroscopy, implying that the polycation is not displaced by cationic liposomes in the formation process. Hence, the polycation-DNA cores (polyplexes) and liposome shells must be oppositely charged to associate. Furthermore, we prove that the liposome:polyplex number ratio (*ρ*_*N*_) is the primary critical parameter to predict stable LPNP formation. We establish that *ρ*_*N*_ ≥ 1 is required to ensure that every polyplex is enveloped by a liposome, avoiding the coexistence of oppositely charged species and thereby inhibiting aggregation. We expect our observations to be valid for the formation of many other LPNPs and composite nanomaterials.

## 1. INTRODUCTION

Non-viral gene delivery to treat genetic and acquired diseases has become clinical reality through the introduction of an siRNA-based treatment for hereditary transthyretin-mediated amyloidosis^1^ and lipid-based mRNA vaccines to address the COVID-19 pandemic^2,3^. Further applications in the pipeline encompass the development of personalized cancer vaccines^4–6^ and permanent knockout of defective genes using CRISPR/Cas9 technology^7^. Most nonviral gene formulations reaching the clinic rely on the use of ionizable cationic lipid formulations as nanocarriers. These materials readily associate with the oppositely charged nucleic acids (NAs), embedding them into internally structured nanoparticles that can be therapeutically efficient *in vivo*^1,8–12^. Notwithstanding this success, the efficacy of non-viral gene therapies remains limited in many cases, particularly for therapeutic targets beyond the liver and immune cells^13^. A promising approach to endow nanocarriers with higher functionality to overcome the many physiological barriers to gene delivery^14–16^ consists in the use of hybrid lipid-polycation-nucleic acid nanoparticles (LPNPs)^17–25^. These nanomaterials consist of a core composed by polycations complexed with NAs (i.e. polyplexes^26^), enveloped by a lipid membrane (Figure 1a). Since the polyplex core and shell components can be independently designed, functionality can be improved^17^, for example through an outer lipid membrane designed for cell-specific targeting and a polycation core optimized for endosomal escape^27^.

**Figure 1.**
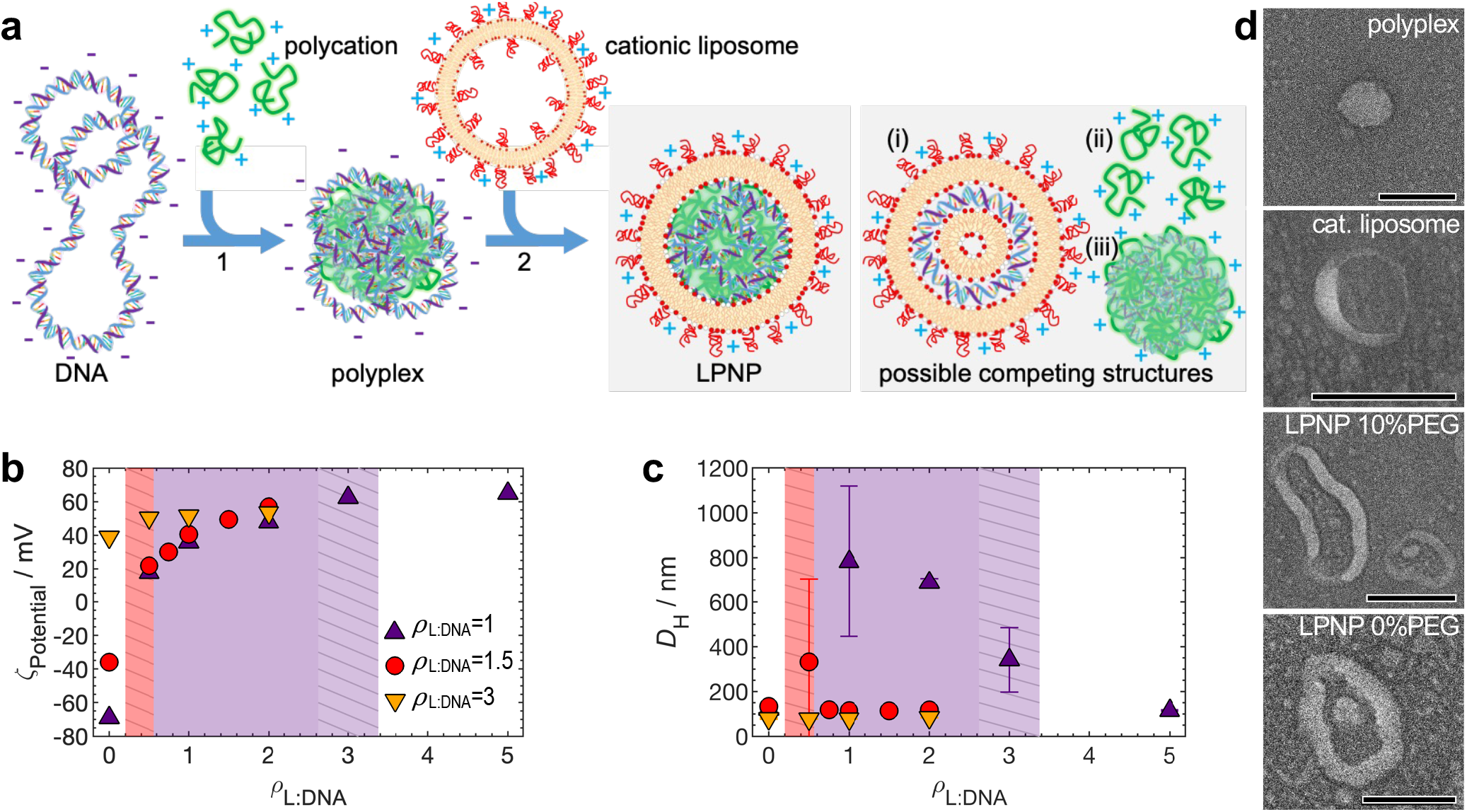
Lipid-Polymer hybrid nanoparticle (LPNP) assembly and physical properties. (**a**) Schematic illustration of the LPNP assembly strategy, consisting of a preliminary step (1) where DNA and polycations are mixed to form polyplexes with an excess negative charge, and a second step (2) where polyplexes are mixed with oppositely charged cationic liposomes to form LPNPs. Along with (or instead of) LPNP formation, hypothetical competing structures can form, such as lipid-DNA complexes (i), polycations released from the polyplex core (ii) or positively charged polyplexes striped from the outer DNA layers (iii). (**b**) ζ potential and (**c**) hydrodynamic diameter (*D*_H_) of the three LPNP systems as a function of *ρ*_L: DNA_. The red dashed area represents the region of the *ρ*_P: DNA_=1.5 system where aggregation was observed in some samples. The purple area represents the region of the *ρ*_P: DNA_=1 system where large aggregation was observed, and the purple dashed area represents the region where aggregation was observed only in some samples. Data are Means ± SD (n=3). (**d**) TEM images of polyplexes, liposomes and LPNPs with 10 and 0 mol% PEG.

LPNPs are complex to produce and characterize since multiple pairwise interactions need to be considered: polycation-DNA, liposome-DNA and liposome-polycation. Typically, the core polyplex is formed first, followed by the addition of oppositely charged liposomes that drive electrostatic complexation between both species, reminiscent of electrostatic-based layer-by-layer assembly^28^. Alternative structures to core-shell LPNPs may form when the polycation and cationic lipids are added to the nucleic acids simultaneously^29,30^, but even when added sequentially, competing interactions between the tree components can still drive the formation of other structures and displace one of the components (Figure 1a).

Elucidating the critical interactions in the LPNP formation process is challenging with widely applied techniques such as Dynamic Light Scattering (DLS), Electrophoretic Mobility, and Transmission Electron Microscopy (TEM). We add dual-colour Fluorescence Cross-Correlation Spectroscopy (FCCS)^31–33^ to this toolset to quantitatively assess the extent of association between polyplexes and liposomes in the formation of LPNPs. The incorporation of FCCS in the standard characterization toolset of other composite materials involving hierarchical assembly is likely to bring further innovation.

## 2. RESULTS AND DISCUSSION

### 2.1. Size and *ζ*_Potential_ reveal stability regions and hint at LPNP formation

A series of samples with varying ratios between lipids, polycations and DNA was prepared to study LPNP assembly behaviour. Sample compositions are defined through the molar ratios of cationic lipids in the liposomes (*ρ*_L:DNA_) and ionizable-cationic amine groups in the polycations (*ρ*_P:DNA_), to anionic groups in DNA. Polylysine, a common cationic polymer in both polyplex and LPNP systems^34,35^, was chosen as polycation, and its concentration fixed at 7 μg/ml. A lipid mixture of DOTAP:DOPC:PEG2K was used for liposome preparation, whereby the cationic lipid DOTAP was maintained in a fixed amount (80 mol%) alongside varying amounts of PEG-lipid.

The methodology for assembling LPNPs is shown in Figure 1a. A polyplex core is assembled in a preliminary step by mixing polylysine with DNA at a fixed *ρ*_P:DNA_ ratio of 1, 1.5 and 3. Subsequently, cationic liposomes are added to form core-shell LPNPs. The isoelectric point (*ρ*_iso_) between polylysine and DNA was determined to be 1.6^36^. *ρ*_P:DNA_ = 1 polyplexes are negatively overcharged with DNA, showing a negative *ζ*_Potential_ (Figure 1b) and are likely to also contain a significant fraction of non-complexed DNA in excess^34^. *ρ*_P:DNA_ = 1.5 polyplexes are close to the *ρ*_iso_, showing a smaller negative *ζ*_Potential_ and are likely to contain just a minute amount of non-complexed DNA, if any. Conversely, *ρ*_P:DNA_ = 3 polyplexes are cationic, and virtually all the DNA is expected to be complexed, coexisting with excess polycations. The size of the polyplexes varies with *ρ*_P:DNA_, being all smaller than 140 nm (Figure 1c). The cationic liposomes presented a hydrodynamic diameter (*D*_*H*_) of ca. 100 nm and high *ζ*_Potential_ values (Table S2 of the supplementary information, SI). The *ζ*_Potential_ and size results for the LPNP formulations are shown in figure 1b-1c. The complete set of data including polydispersity index (PDI) can be found in Table S3.

Figure 1b-1c indicates two trends for the LPNP systems depending on the surface charge of the starting polyplex. For samples obtained from cationic polyplexes (*ρ*_P:DNA_ = 3) neither size nor *ζ*_Potential_ are significantly affected when cationic liposomes are added. Conversely, when the starting polyplex presents negative *ζ*_Potential_ (*ρ*_P:DNA_ = 1 and 1.5), LPNP charge switches to cationic already at the lowest *ρ*_L:DNA_ additions, coinciding with the observation of strong aggregation (*D*_*H*_ > 500nm), especially for *ρ*_P:DNA_ = 1. Further addition of liposomes leads to stabilization. Clear solutions with *D*_*H*_ < 120nm are obtained for *ρ*_L:DNA_ = 5 and *ρ*_L:DNA_ = 0.75 for *ρ*_P:DNA_ = 1 and *ρ*_P:DNA_ = 1.5 LPNPs, respectively. Since *ρ*_P:DNA_ = 1 polyplexes are more negatively charged and are expected to coexist with significant amounts of free DNA in solution (which needs to be neutralized), more liposomes are needed to stabilize these LPNPs when compared with the *ρ*_P:DNA_ = 1.5 system. Yet, at a first glance, the factor of ca. six times larger *ρ*_L:DNA_ needed to stabilize *ρ*_P:DNA_ = 1 polyplexes seems high. TEM results (Figure 1d) confirm the formation of LPNPs, with imaged particles showing a core like the polyplex samples, enveloped by a membrane.

Negative polyplexes aggregating initially and then redispersing for increasing *ρ*_L:DNA_ hints at strong electrostatic interactions with liposomes and conversion of polyplexes into LPNPs. One should however also consider other interactions potentially leading to the formation of structures competing with LPNPs (Figure 1a). For instance, DNA could be partially or entirely displaced from the polycation to the cationic liposomes to form lipoplexes. Such structures could have sizes and *ζ*_Potential_ like LPNPs. Regarding the positively charged polyplexes (*ρ*_P:DNA_ = 3), the lack of observable changes with *ρ*_L:DNA_ could hint at no formation of LPNPs, but also here one should consider the possibility of cationic liposomes displacing some polycations to form LPNPs. These alternative scenarios are difficult to discern with DLS/electrophoretic mobility and even with TEM.

### 2.2. Dual-colour fluorescence time traces provide semi-quantitative colocalization between cationic liposomes and polyplexes

Colocalization between two entities labelled with two distinct fluorophores can be assessed qualitatively by looking at the spatial overlap between the two labels using fluorescence microscopy^22^. However, this approach is challenging to render quantitative^37^. FCCS can overcome this limitation and provide a quantitative measure of colocalization along with simultaneous determination of the sizes of the labelled species^31,32^. FCCS has been successfully used to characterize interactions between biologic species^31,32,38,39^, and to assess the formation^33,40^ (and disassembly^41^) of cationic liposome-DNA complexes, polymeric^42^ and silica^43,44^ nanoparticles, and liposome-coated metal organic frameworks^45^.

To monitor the formation of LPNPs (Figure 1a), we label polylysine covalently with Atto-488 and likewise the cationic liposomes with Texas Red. Since both labelled species are cationic, they can only associate if DNA acts as an intermediary. Hence, colocalization between polylysine and liposomes indicates association between the liposomes and polyplex cores. Simultaneously, this rules out other competing structures in which DNA would be displaced, as illustrated in Figure 1a. If DNA is labelled instead of polycations, the absence of cross-correlation between both signals still discards the presence of LPNPs, but its presence does not allow for the distinction between LPNPs and other competing structures (e.g., lipoplexes). In the case of formulations starting with cationic polyplexes (*ρ*_P:DNA_ = 3), the existence of labelled polycation free in solution makes the FCCS measurements difficult to analyse, and in this case the initial measurements were complemented with measurements whereby the DNA is labelled (YOYO-1, green) instead of the polycation.

In FCCS, the confocal volumes defined by the green and red lasers of a confocal microscope are fixed in space, while labelled species diffuse in and out of the overlapping green and red region (Figure 2a). If polyplexes and liposomes do not form LPNPs, they diffuse independently, producing uncorrelated spikes in the two fluorescence signals. This is the case for formulations starting with *ρ*_P:DNA_ = 3 polyplexes, in which polyplexes and cationic liposomes show unsynchronized spikes in the fluorescence channels (Figure 2b). In contrast, if polyplexes and liposomes form LPNPs, they diffuse together producing correlated fluorescence signals. This is the case for the formulations starting with *ρ*_P:DNA_ = 1.5 polyplexes (Figure 2c), where excellent overlap between the green and red signals is observed. For formulations starting with *ρ*_P:DNA_ = 1 (Figure 2d), the spikes in the green channel overlap with red spikes, but there are many red spikes that have no overlap with green. This is because a high excess of liposomes is present, producing only red spikes. Overall, even if somewhat qualitative, these observations confirm LPNP formation when the charge is being inverted (*ρ*_P:DNA_ = 1 and 1.5), but not when the charge of polyplexes and liposomes has the same sign (*ρ*_P:DNA_ = 3).

**Figure 2.**
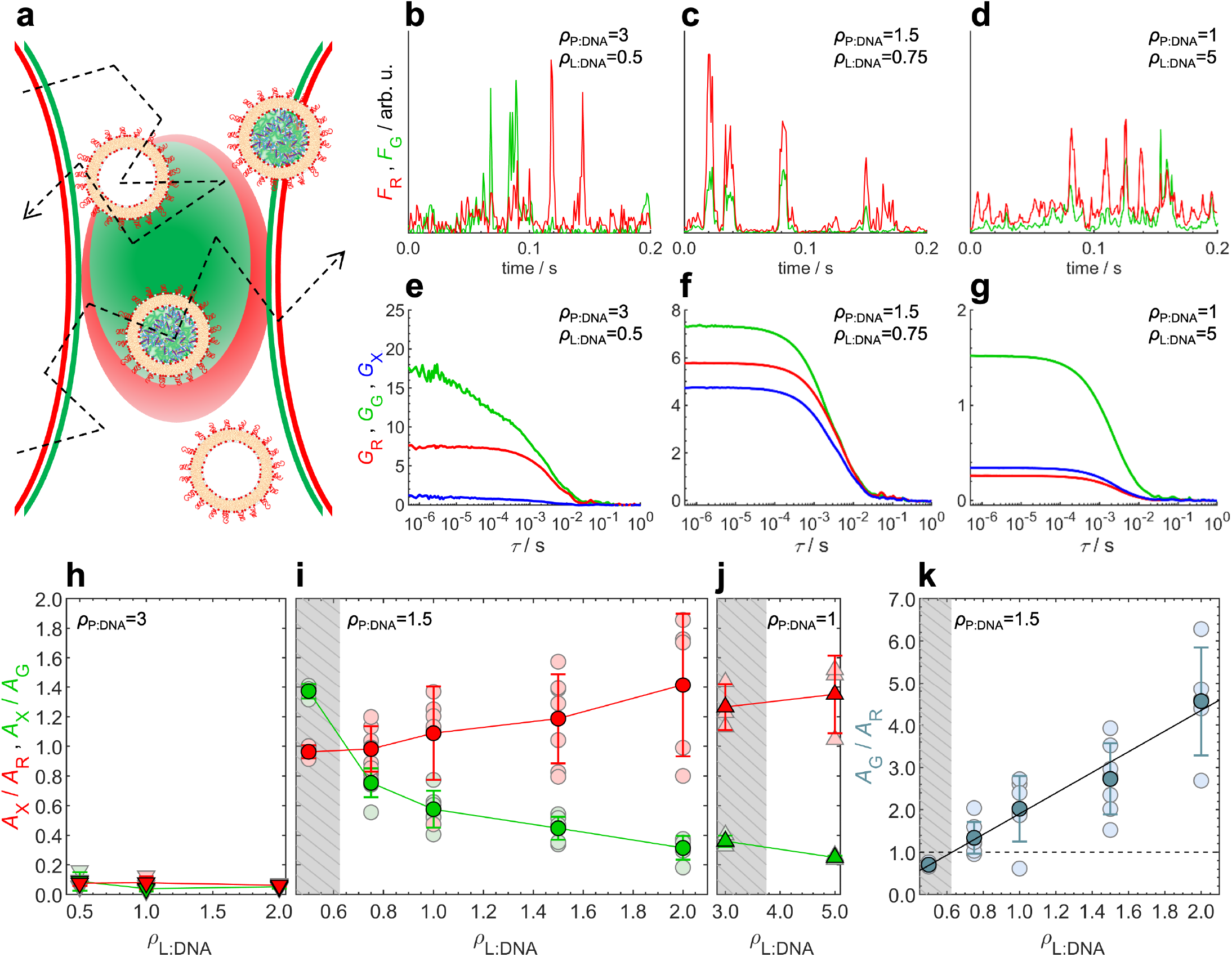
FCCS quantifies association between cationic liposomes and polyplexes. (**a**) Schematic illustration of the green and red confocal volumes with LPNPs and cationic liposomes diffusing through them. Panels **b**-**d** show representative fluorescence time traces for samples *ρ*_P: DNA_=3, *ρ*_L: DNA_=0.5; *ρ*_P: DNA_=1.5, *ρ*_L: DNA_=0.75; and *ρ*_P: DNA_=1, *ρ*_L: DNA_=5, respectively. Panels **e**-**g** show the corresponding auto and cross-correlation curves for the same samples. The green and red fluorescence (*F*_G_ and *F*_R_) and auto-correlation curves (*G*_G_ and *G*_R_) are shown in green and red, respectively. The cross-correlation curves (*G*_X. meas_) are shown in blue. Non-associating particles, like cationic polyplexes and cationic liposomes, diffuse independently, producing uncorrelated green and red fluorescence fluctuations, visible both in the time traces (**b**) and in the negligeable amplitude of *G*_X. meas_. in (**e**). LPNPs, containing both a polyplex (labelled with Atto488, green) and a cationic liposome (Texas-Red) produce correlated green and red fluorescence fluctuations, visible both in the time traces (**c** and **d**), and large amplitudes of *G*_X. meas_ (**f** and **g**). Panels **h**-**j**, show the *A*_X_/*A*_R_ (red symbols) and *A*_X_/*A*_G_ (green symbols) ratios, which provide the fraction of polyplexes converted to LPNPs and the fraction of liposomes used in LPNPs, respectively. Individual measurements are represented by light-red and light-green symbols, while their means and respective error bars are represented by dark-red and dark-green. *A*_X_/*A*_R_ and *A*_X_/*A*_G_ are very low for the *ρ*_P: DNA_=3 system, confirming negligeable association between polyplexes and liposomes with the same charge sign (**h**). The *A*_X_/*A*_R_ ratio for *ρ*_P: DNA_=1.5 (**i**) and *ρ*_P: DNA_=1 (**j**) systems is ≈1 for low *ρ*_L: DNA_ and increases slightly as *ρ*_L: DNA_ increases, indicating full conversion of polyplexes to LPNPs, and an average number of liposomes per LPNP increasing slightly beyond one with *ρ*_L: DNA_. Conversely, *A*_X_/*A*_G_ decreases steadily with *ρ*_L: DNA_, indicating that after colloidal stability is reached, most added liposomes are in excess, not associating with LPNPs. Panel **k** shows the *A*_G_/*A*_R_ ratio (for *ρ*_P: DNA_=1.5), which is equivalent to the liposome:polyplex ratio (*ρ*_N_). The region of robust colloidal stability for LPNPs shows *ρ*_N_>1. The region of marginal stability shows *ρ*_N_<1. This suggests that *ρ*_N_≈1 is a requirement for robust stability. The boundary between marginal and robust stability (*ρ*_N_=1) is estimated at *ρ*_L: DNA_=0.63 for *ρ*_P: DNA_=1.5. The grey-dashed areas in **h**-**k** delineate the compositions of marginal stability, in which some samples were not measurable due to the presence of aggregates. Data are Means ± SD (n≥3).

### 2.3. FCCS auto- and cross-correlation quantify association between cationic liposomes and polyplexes

For a quantitative analysis, the auto- and cross-correlation curves (resulting from time traces) can be examined^31,33^ (Figures 2e-g). The green and red auto-correlation curves describe the dynamics of species carrying the green and red labels, respectively. Importantly, the cross-correlation curve results from the correlation between the two signals and therefore only contains information on the species carrying both labels. The decay time constants of these curves are related with the species’ diffusion coefficients (Eqs. S2-S3) and are used to extract LPNP size (Table S4 and Figure S1). The amplitudes of the correlation curves contain information on the number of each labelled specie, and on the number of species containing both labels^31,37^. Hence, the fraction of LPNP formation can be determined. For the simplified case of: (i) one polyplex associating with one cationic liposome (1:1 stoichiometry) to form one LPNP; (ii) no crosstalk between the two channels; and (iii) no changes in the brightness of the species following the association, the equations relating the amplitudes of the correlation curves with the number of species in the confocal volume become straightforward (Eqs 1a-c).

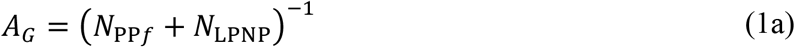

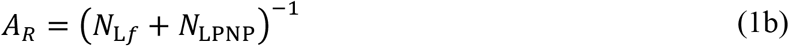

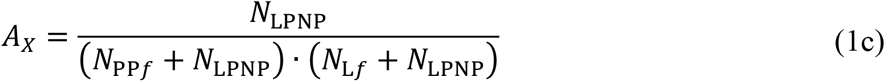

Here, *A*_*G*_ and *A*_*R*_ are the amplitudes of the green and red auto-correlation, respectively, and *A*_*X*_ is the amplitude of the cross-correlation. *N*_*PPf*_, *N*_*Lf*_, and *N*_*LPNP*_, are the average number of free polyplexes, free liposomes and LPNPs in the confocal volumes, respectively. The ratio between *A*_*X*_ and *A*_*R*_ provides a direct measure of the fraction of polyplexes that were converted to LPNPs (*f*_*LPNP*_), according to:

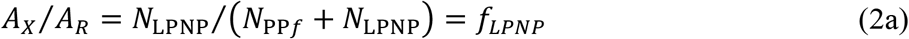

Conversely, the *A*_*X*_/*A*_*G*_ ratio provides a direct measure of the fraction of cationic liposomes used to make LPNPs:

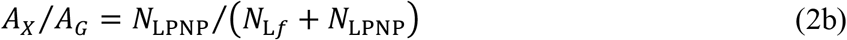

The approximations above are reasonable since the experimental conditions are such that the crosstalk from the green to the red channel is kept low at ca. 10%, allowing for its correction^46^, and the brightness of polyplexes and liposomes do not change significantly upon association. In section S2 of the SI we detail why the 1:1 stoichiometry assumption is appropriate to estimate *f*_*LPNP*_ and a much more complex model based on 1:*n* stoichiometry with *n* liposomes per LPNP is not required.

The overlap volume correction factor (*V*_*crct*_ = 74.6%), is determined in a controlled experiment using mixtures of single-labelled and double-labelled PEGylated liposomes (section S3 and Figure S3)^47^. After correcting the crosstalk^46^, the *A*_*X*_ used in Eqs. 1-2 is obtained from the measured amplitude of the cross-correlation (*A*_*X*.*meas*_) and *V*_*crct*_ according to:

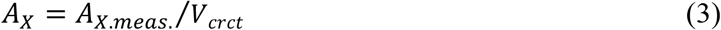

### 2.4. Liposomes and polyplexes with the same charge do not associate

Starting with the *ρ*_P:DNA_ = 3 system, where we had seen low synchronization between the green and red signals, Figure 2e shows that the amplitude of the cross-correlation curve *A*_*X*.*meas*_. is much lower than the amplitudes of the autocorrelation curves, resulting in *A*_*X*_/*A*_*R*_ and *A*_*X*_/*A*_*G*_ ratios very close to zero (Figure 2h). As mentioned above, in this *ρ*_P:DNA_ = 3 case, the DNA was labelled instead of the polycation, but the absence of cross-correlation still proves lack of association between polyplexes and liposomes and virtually no LPNP formation. The correlation curves with polycations labelled instead are shown in Figure S4.

Like-charged polyplexes and liposomes are unlikely to associate unless there is a strong non-electrostatic interaction between them, or the liposomes replace part of the polycation in the polyplexes. Notwithstanding, some systems composed by cationic polyplexes and cationic liposomes have reported better transfection than the polyplexes alone^48^. It is unclear if in such cases the use of weaker polycations (e.g., polyethyleneimine – PEI) can lead to partial replacement of the polymer by liposomes to form LPNPs or if the presence of liposomes can boost transfection of polyplexes even without LPNP formation.

### 2.5. Oppositely charged liposomes and polyplexes near *ρ*_iso_ associate strongly

In contrast with cationic polyplexes, the *ρ*_P:DNA_ = 1.5 system shows a much larger *A*_*X*.*meas*_., comparable to both *A*_*G*_ and *A*_*R*_, indicating strong colocalization (Figure 2f). Since in this case, both labelled species are cationic (polycation with green and liposomes with red), their colocalization indicates strong association between polyplexes and liposomes.

Figure 2i, shows the corrected *A*_*X*_/*A*_*R*_ and *A*_*X*_/*A*_*G*_ ratios as a function of *ρ*_L:DNA_. It is noticeable that while the mean values present well-defined and robust trends, the individual measurements (typically five-six independent samples) are significantly noisier, especially *A*_*X*_/*A*_*R*_. This occurs because FCCS operates in a quasi-single-particle regime, requiring long measurement times or multiple measurements to reach adequate statistics. Since long measurements are not feasible due to sample drying, shorter measurements were performed with several independently prepared samples, exhibiting a larger variability that vanishes when computing the mean. Note also that while the variability of *A*_*X*_/*A*_*R*_ is significant, the variability of *A*_*X*_/*A*_*G*_ is much lower. This is expected if most polyplexes are converted to LPNPs coexisting with excess liposomes. The apparently noisier nature of the individual measurements is therefore not of concern, as can be also confirmed by the results of LPNPs assembled with 0 and 5% PEG liposomes, which show the same trends—noisy individual *A*_*X*_/*A*_*R*_ measurements, but robust means with well-defined trends (Figure 3a).

**Figure 3.**
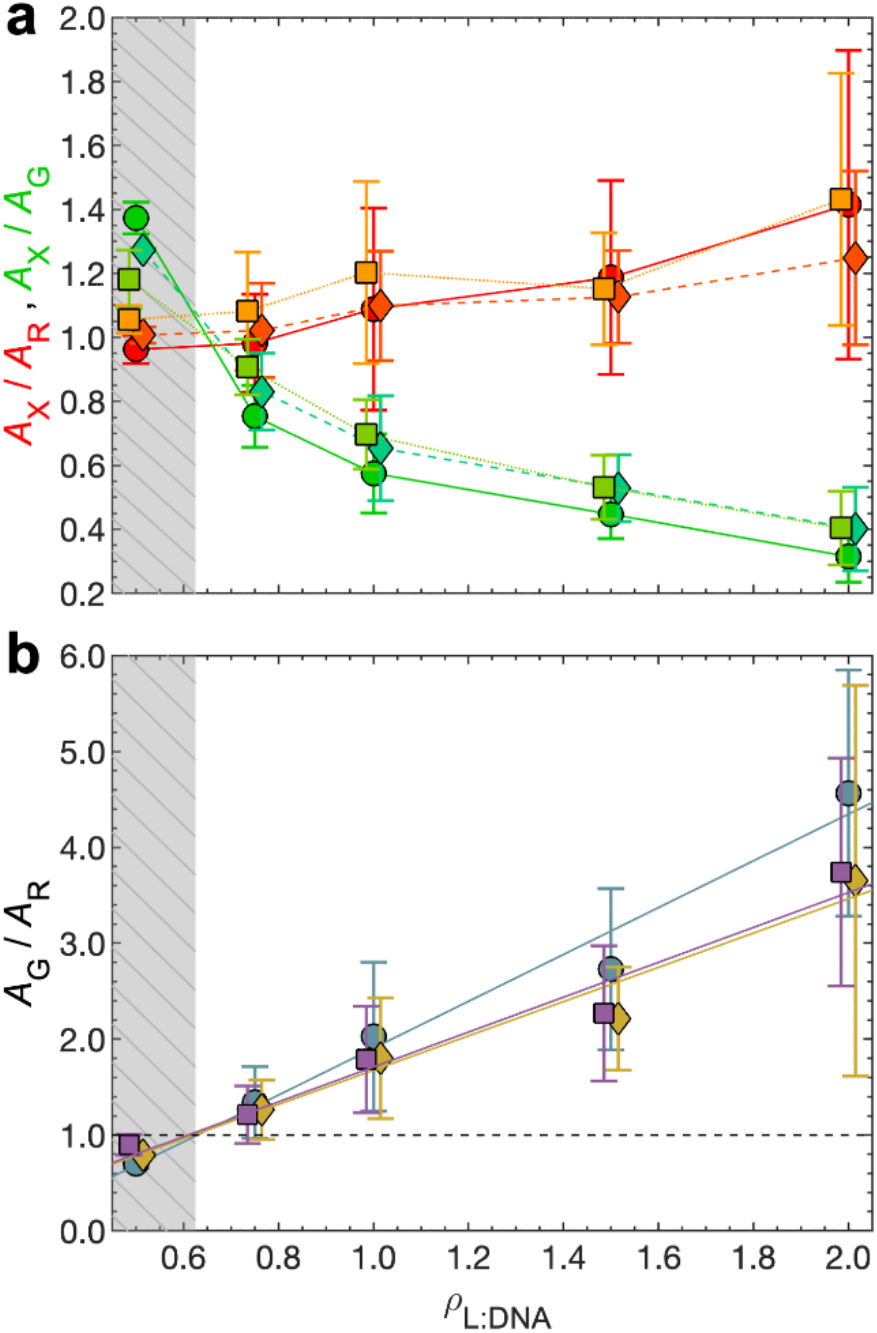
Effect of liposome PEGylation in the assembly of LPNPs. (**a**) *A*_X_/*A*_R_ (red-orange symbols) and *A*_X_/*A*_G_ (green-blue symbols) ratios for LPNP systems with 0 mol% PEG (squares), 5 mol% PEG (diamonds) and 10 mol% PEG (circles) liposomes. The trends are similar in the three systems, but there is a systematic small increase of the *A*_X_/*A*_R_ and *A*_X_/*A*_G_ pair for lower degrees of liposome PEGylation. This indicates that the conversion from polyplexes to LPNPs is practically complete for all three systems, but the fraction of used liposomes and average number of liposomes per LPNP increase slightly for lower degrees of liposome PEGylation. Note that similarly to Figures 2i-j, also here the *A*_X_/*A*_R_ error bars are large, due to the quasi-single-particle nature of the measurements, but the mean values are robust and follow coherent trends. (**k**) *A*_G_/*A*_R_ ratio for LPNP systems with 0 mol% PEG (squares), 5 mol% PEG (diamonds) and 10 mol% PEG (circles) liposomes. The trends are identical for the three systems. The regions of robust colloidal stability for LPNP formation show *ρ*_N_>1, whereas the regions of marginal stability show *ρ*_N_<1. The boundaries between marginal and robust stability (*ρ*_N_=1) are the same for the three systems (*ρ*_L: DNA_=0.63). This reinforces that *ρ*_N_≥1 is a requirement for robust stability. The fact that the linear fits to *A*_G_/*A*_R_ have a lower slope for the 0 and 5 mol% PEG systems is caused by the greater deviations to the 1:1 stoichiometry in these systems. Data in **a** and **b** is slightly offset in the horizontal axis for easier visualization. The grey-dashed areas delineate the compositions of marginal stability, in which some samples were not measurable due to the presence of aggregates. Data are Means ± SD (n≥3).

Within the 1:1 liposome:polyplex stoichiometry ratio approximation *A*_*X*_/*A*_*R*_ is equal to *f*_*LPNP*_ (Eq 2a). It is noticeable that *A*_*X*_/*A*_*R*_ increases from ca. 1 to 1.42 as *ρ*_L:DNA_ is increased from 0.5 to 2. This indicates a practically full conversion of polyplexes into LPNPs (*f*_*LPNP*_≈1) in detriment of the alternative structures depicted in Figure 1a. The fact that the average *A*_*X*_/*A*_*R*_ values become larger than one for higher *ρ*_L:DNA_ indicates that for larger amounts of added liposomes, some LPNPs may have two liposomes attached, resulting in a 1:*n* stoichiometry with *n* slightly greater than 1, but still with *f*_*LPNP*_≈1. A more detailed analysis considering the 1:*n* stoichiometry is provided in section S2, confirming *f*_*LPNP*_≈1. Under such conditions *n*∼1.7 for *ρ*_L:DNA_ = 2, which indicates that even at the largest *ρ*_L:DNA_, most LPNPs will be associated with either one or two liposomes, and most liposomes added will be in excess.

Figure 2i also shows the *A*_*X*_/*A*_*G*_ ratio, which directly provides the fraction of liposomes used to make LPNPs (Eqs 3b). *A*_*X*_/*A*_*G*_ is greater than one for *ρ*_L:DNA_ = 0.5, which suggests that for the smallest *ρ*_L:DNA_ there may not be enough liposomes to completely coat the polyplexes. This helps explaining why in this composition LPNP solutions are sometimes colloidally unstable. As *ρ*_L:DNA_ is increased, *A*_*X*_/*A*_*G*_ starts to steadily decrease, indicating that there is now an excess of liposomes in solution that do not associate with the polyplexes. This supports a 1:*n* stoichiometry in which *n* is relatively close to one.

### 2.6. FCCS reveals that the liposome:polyplex number ratio *ρ*_N_ is the critical parameter determining stable LPNP formation, not the charge ratio

In the previous section it became apparent that the ratio between the number of liposomes and polyplexes (*ρ*_*N*_) is important. This quantity is not straightforward to obtain with most techniques but is easily accessible using FCCS. The total number of polyplexes and liposomes in solution is obtained through 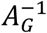 and 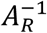, respectively (Eqs 1a-b). If *n*≈1, the *A*_*G*_/*A*_*R*_ ratio then provides *ρ*_*N*_. (Since *n*≈1.7 mostly when there are also many free liposomes, this results in a small error.)

Figure 2k shows that for the *ρ*_P:DNA_ = 1.5 system, all the samples in the stable region (*ρ*_L:DNA_ ≥ 0.75) have a *ρ*_*N*_ ratio greater than one. Conversely, the samples in the marginally stable region (*ρ*_L:DNA_ = 0.5) show a *ρ*_*N*_ slightly below one. This provides a very strong hint that *ρ*_*N*_ ≥ 1 is a critical requirement to obtain robust colloidal stability.

If the number of liposomes is smaller than the number of polyplexes, some polyplexes will be transformed into cationic LPNPs while some others will remain anionic. Hence, even in a situation of large overall positive charge, if *ρ*_*N*_ < 1 the interactions between non-converted anionic polyplexes and converted cationic LPNPs can lead to aggregation. Provided that each liposome has an absolute charge greater than the polyplexes, and no excess DNA present, *ρ*_*N*_ ≥ 1 ensures that all polyplexes are converted, and LPNPs are stable.

However intuitive, this *ρ*_*N*_ ≥ 1 requirement is a critical insight. The reason why we and others have failed to think of it is because *ρ*_*N*_ is an elusive quantity, difficult to estimate beforehand or determine experimentally. Furthermore, it depends on sample preparation. Hence, *ρ*_*N*_ is not a salient feature in sample preparation and one tends to focus on absolute charge ratios instead. The latter are easier to control experimentally, but less useful in predicting the association between liposomes and polyplexes, since absolute charges are distributed over a discrete number of particles. FCCS then plays a crucial role in revealing and highlighting *ρ*_*N*_ by measuring both the number of polyplexes and liposomes simultaneously.

### 2.7. Polyplexes far from *ρ*_iso_ are more numerous, requiring more liposomes to form LPNPs

Overall, *ρ*_P:DNA_ = 1 polyplexes show similar trends to the *ρ*_P:DNA_ = 1.5 system (Figures 2g and 2j). Namely, all the polyplexes are converted into LPNPs (*f*_*LPNP*_≈1) and the average number of liposomes per LPNP (*n*) is also slightly greater than one, as seen from the *A*_*X*_/*A*_*R*_ ratio slightly above one. However, there are two important additional points to highlight.

The first is that when significantly away from *ρ*_iso_, some of the DNA is expected to remain free in solution, non-complexed^34^. This free DNA also needs to be complexed by the cationic liposomes for the system to become colloidally stable, thus requiring a larger *ρ*_L:DNA_ ratio. Liposomes complexed with this excess DNA are not expected to be associated with polyplexes, which explains the observation that *A*_*X*_/*A*_*G*_ is measured at ca. 0.25. However, contrary to the *ρ*_P:DNA_ = 1.5 case seen above, where a decreasing *A*_*X*_/*A*_*G*_ is just indicative of superfluous liposomes, here, these extra liposomes are needed to neutralize free DNA, and almost certainly this indicates coexistence of LPNPs with lipoplexes. This partially explains the much larger *ρ*_L:DNA_ ratio needed to obtain stability in the *ρ*_P:DNA_ = 1 systems.

The second is that the number of polyplexes is significantly greater for *ρ*_P:DNA_ = 1 than *ρ*_P:DNA_ = 1.5. This can be seen by the amplitudes of the green autocorrelation curves in Figures 2f and 2g (∼5x greater – recall Eq 2a) and Table S5 (∼3.5x greater). The greatest consequence of this larger number of polyplexes is that to ensure the *ρ*_*N*_ ≥ 1 requirement, the number of liposomes needed to stabilize the system also increases. This, together with the excess of DNA discussed above, helps justifying the much higher *ρ*_L:DNA_ needed for *ρ*_P:DNA_ = 1 LPNPs.

### 2.8. The amount of PEG on liposomes influences the number of liposomes per LPNP

The findings above are largely reproduced when the 10 mol% PEG liposomes are replaced by 5 and 0 mol% PEG liposomes (Figure 3a). Both *A*_*X*_/*A*_*R*_ and *A*_*X*_/*A*_*G*_ ratios show similar trends across the three systems, indicating *f*_*LPNP*_≈1 and *n* increasing slightly beyond one for large *ρ*_L:DNA_. However, it is also visible that as the PEG amount in the liposomes decreases, both the *A*_*X*_/*A*_*R*_ and *A*_*X*_/*A*_*G*_ ratios increase. This is more evident for *A*_*X*_/*A*_*G*_, since the *A*_*X*_/*A*_*R*_ values have larger error bars. Overall, both these ratios indicate that while *n* is expected to still be relatively small, it increases when the PEGylation degree decreases. Since the PEG coating on liposomes acts as a steric barrier against adsorption, it is not surprising that on average low-PEG liposomes complex more extensively with polyplexes.

Regarding the *ρ*_*N*_ ratio, which is approximated by *A*_*G*_/*A*_*R*_ (Figure 3b), also here we find *ρ*_*N*_ > 1 for all stable LPNPs, and *ρ*_*N*_ < 1 for the non-stable ones, which reinforces that *ρ*_*N*_ ≥ 1 is a critical requirement for LPNP colloidal stability. The boundary between marginal and robust stability (*ρ*_*N*_ = 1) is similar for the three systems.

### 2.9. General features of LPNP assembly behaviour

Figure 4 summarises the results and observations of this work in the form of a ternary map of assembly in excess water (given the out-of-equilibrium nature of the system, the map is not strictly a phase diagram). Additional details regarding the boundaries on the map and its implications are available in section S4.

**Figure 4.**
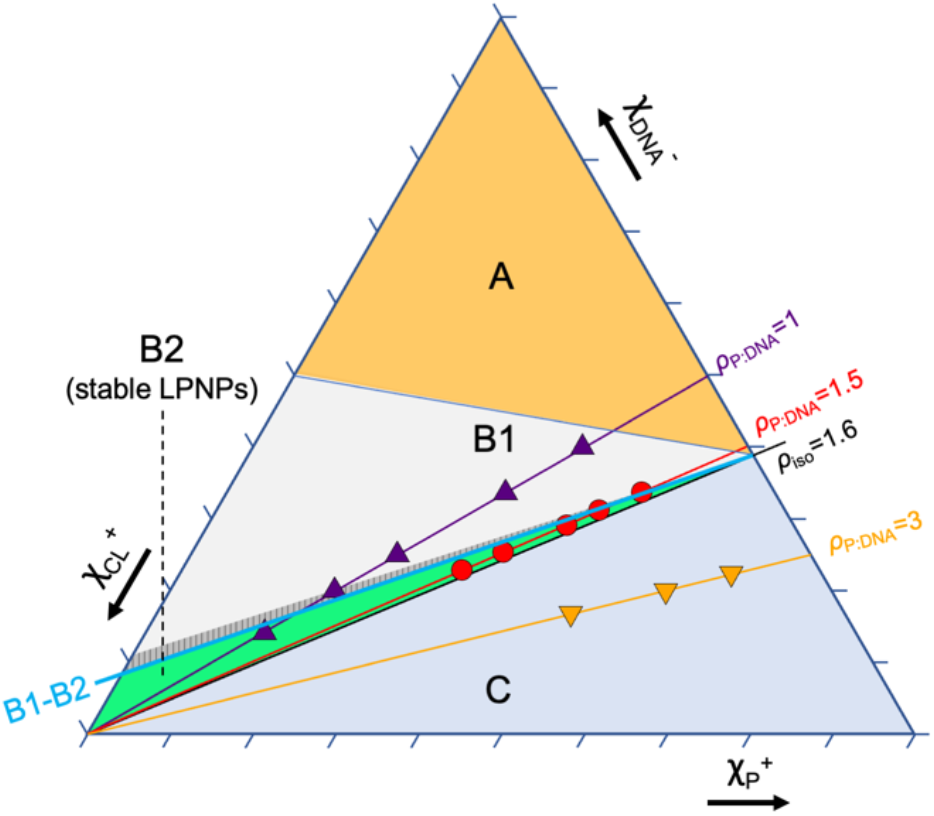
Ternary lipid-polylysine-DNA assembly map in excess water. The three axes represent the charge molar fractions of DNA 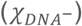, polylysine 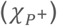 and cationic liposomes 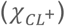. The polylysine concentration is fixed at 7 μg/mL. The map is subdivided in three main regions. Region A encompasses overall negatively charged systems (not investigated). In region B the starting polyplexes are negative but the overall charge is net-positive via addition of cationic liposomes. In sub-region B1 the assemblies are not colloidally-stable whereas in B2 LPNPs stable for at least one week are formed. The B1-B2 boundary between aggregation and stable LPNP formation is set by the liposome:polyplex ratio (*ρ*_N_), which must be *ρ*_N_ ≥1 when there is no excess DNA present (see text). The patterned area immediately above the B1-B2 boundary indicates the region where colloidal stability is observed only in some samples. The number of liposomes and polyplexes (hence *ρ*_N_) is defined by both the composition and methodology. Within B2, *ρ*_P: DNA_=1.5 LPNPs coexist with excess cationic liposomes, whereas *ρ*_P: DNA_=1 LPNPs coexist with lipoplexes. In region C the starting polyplexes are positive (below the *ρ*_iso_ line), and no LPNP formation was observed after addition of cationic liposomes.

The first important observation is the lack of formation of LPNPs when starting with positive polyplexes (region C on the map). This indicates that the interaction between polylysine and DNA is robust, with the polycation not being displaced by liposomes. The second important observation is the need to have at least one liposome per polyplex (*ρ*_*N*_ ≥ 1) to form stable LPNPs. This ensures that every polyplex is enveloped by a liposome, having their negative charge reversed and inhibiting aggregation. This requirement divides region B into two subregions: B1, for *ρ*_*N*_ < 1, where aggregation is observed, and B2, for *ρ*_*N*_ ≥ 1, where LPNPs stable for at least one week are formed. In B2, the conversion of polyplexes to LPNPs is complete, indicating that neither the polycation nor DNA are displaced by the liposomes.

While the location of the B1-B2 boundary is specific to this system, we postulate that the general features of LPNP assembly observed in this work can be transposed to systems containing reasonably strong polyelectrolytes and cationic liposomes with monovalent lipids and similar membrane rigidity. If weak polycations (with low ionization degree) are used in the formation of the polyplex cores, or multivalent cationic liposomes as the shell, enhanced lipid-DNA interactions in detriment of polycation-DNA could potentially lead to displacement of the polycation and modify the assembly behaviour. Regardless, the FCCS analysis approach detailed here can be extended to quantify the association between different components in complex systems and provide valuable insights into the overall assembly behaviour of LPNPs and other composite materials for diverse applications besides gene delivery.

## Acknowledgements

We are grateful to Cyrus Safinya (UCSB), Ulf Olsson (Lund U.) and Paulo Freitas (INL) for useful discussions. This research is supported by the Microfluidic Layer-by-layer Assembly of Cationic Liposome - Nucleic Acid Nanoparticles for Gene Delivery project (032520) co-funded by FCT and ERDF through COMPETE2020. JLP is supported by a post-doctoral fellowship from the Ramón Areces Foundation (reference BEVP30A5827).

## Online content

The supplementary data contains a detailed FCCS implementation description, 1:*n* polyplex:liposome stoichiometry model of LPNP formation, and determination of the overlap confocal volume.

## 3. MATERIALS AND METHODS

### Materials

The lipids 1,2-dioleoyl-3-trimethylammonium-propane (DOTAP), 1,2-dioleoyl-sn-glycero-3-phosphocholine (DOPC) and 1,2-distearoyl-sn-glycero-3-phosphoethanolamine-N-[amino(polyethylene glycol)-2000] (DSPE-PEG) solubilized in chloroform were purchased from Avanti Polar Lipids and used as received. Poly-L-lysine solution 0.1 % (w/v) in water was purchased from Sigma-Aldrich. GFP plasmid (pCMV-GFP), obtained through Addgene, was a gift from Connie Cepko (Addgene plasmid #11153)^49^. The dye Atto 488-NHS was purchased from ATTO-TEC GmbH. Texas Red 1,2-Dihexadecanoyl-snGlycero-3-Phosphoethanolamine, Triethylammonium salt (Texas Red-labelled lipid) and the dye YOYO-1 were purchased from ThermoFisher Scientific Inc. All the aqueous solutions were prepared with DNAse-free and RNAse-free MilliQ water.

### Characterization techniques

The different nanostructures obtained in this work were characterized by the following techniques: DLS and electrophoretic mobility (*ζ*_Potential_) were performed with a SZ-100 device from Horiba, measuring scattering at 173°. TEM of the samples after negative staining with UranyLess© (following the manufacturer’s instructions) was performed in a JEOL 2100 200kV TEM. FCCS was performed in a confocal microscope LSM780 from Zeiss following a previously described protocol^31^, with 5 s per measurement and carrying out 40 repeats per sample (excitation laser lines: 488 nm for Atto 488 dye and 561 nm for Texas Red). Laser power was set so that the intensity in the “red” (561 nm) channel was slightly above that in the “green” (488 nm) channel, thus minimizing overestimation of cross-correlation due to crosstalk^46^. FCCS auto- and cross-correlation curves were fitted to a 3D normal diffusion model^50^ using QuickFit 3.0 software and home-built Matlab scripts. After calibrating the FCCS setup employing fluorophores with a known diffusion coefficient^51^, the fits obtained for the cross-correlation curves were used to obtain the diffusion coefficient of the LPNPs.

### Labelling of polylysine with Atto 488 dye

Polylysine was labelled by reacting the polymer with Atto 488-NHS (ratio 1:100 Atto dye: NH_2_ groups in the polymer). After stirring in aqueous solution at room temperature overnight, the labelled polymer was obtained. The purification was performed by size exclusion chromatography using Sephadex G25® stationary phase.

### Preparation of LPNPs

Polyplexes were prepared by mixing aqueous solutions of plasmidic DNA and polylysine at different charge ratios (CR, ratio between positive charges in the polymer to negative charges in the plasmid: 1, 1.5 and 3), and incubating them at room temperature for 30 min in an orbital shaker.

PEGylated cationic liposomes (composition DOTAP/DOPC/DSPE-PEG 80/10/10) were prepared. First, the lipids were mixed in chloroform solution, followed by evaporation of the solvent, rehydration with water and sonication with a tip sonicator at a final total lipid concentration of 4 mM (sonication conditions: 10% amplitude, 1 minute, 50% duty cycle using a Branson Digital Sonifier® 250 Model). Cationic liposomes formulations with the ratios 80/15/5 and 80/20/0 were also prepared.

Finally, both components (polyplex and liposomes) were mixed at different proportions, with a lipid N/DNA P ratio in the range of 0.5-2. A final polylysine concentration of 7 μg/mL was fixed for all the prepared samples. The obtained dispersions were characterized by *ζ*_Potential_ and DLS.

For FCCS, the LPNPs were prepared following the same protocol but employing labelled components. Atto 488-labeled polylysine was used to prepare the polyplexes, while Texas Red-labelled phospholipid was included in the liposomal formulation (0.2 % total lipid weight). Some samples were also prepared with YOYO 1-labeled DNA and Texas Red-labelled liposomes.

