## Supplementary Information for "Stability Criterion for the Assembly of Hybrid Lipid-Polymer-Nucleic Acid Nanoparticles"

**for**

**S1. Fluorescence Cross-Correlation Spectroscopy background**

In the most common experimental implementation of Fluorescence Cross-Correlation Spectroscopy (FCCS), two components of a system are labelled with two spectrally resolved fluorophores (*a* and *b*, typically emitting in the green and red spectra) and their fluorescence signals ($F_a$ and $F_b$) are followed as they diffuse in and out of the two corresponding confocal volumes. The overlap between the two confocal volumes should be as large as possible. The dynamics of the components cause fluorescence intensity fluctuations that can be represented through the normalized correlation function (*G*), determined from the time-dependent fluorescent signals:

$$G_{ab}(\tau) = \frac{\langle \delta F_a(t) \delta F_b(t+\tau) \rangle}{\langle F_a(t) \rangle \langle F_b(t) \rangle} \quad \text{(S1)}$$

where $\delta F(t) = F(t) - \langle F(t) \rangle$ and $\tau$ is the lag time. Autocorrelation functions use only one signal (*a*=*b*) and represent the dynamics of species containing that signal. The cross-correlation uses both signals (*a*≠*b*) and represents only the dynamics of species containing the two labeled components. FCCS can therefore be used to quantify colocalization between two labeled species. For simplicity the cross-correlation function will be denoted by $G_X$.

In the case of free diffusion in three dimensions, the correlation function *G* can be represented through:

$$G(\tau) = A \cdot M(\tau) \quad \text{(S2)}$$

where the amplitude $A$ contains information on the number of species, and $M(\tau)$, which contains information regarding their dynamics, is given by:

$$M(\tau) = \left(1 + \frac{4D\tau}{w_0^2}\right)^{-1} \left(1 + \frac{4D\tau}{w_z^2}\right)^{-1/2} = \left(1 + \frac{\tau}{\tau_D}\right)^{-1} \left(1 + \frac{\tau}{\tau_D S^2}\right)^{-1/2} \quad \text{(S3)}$$

Here, $S = w_z/w_0$ is the aspect ratio between the axial and lateral radii of the detection volume ($w_z$ and $w_0$, respectively), $\tau_D = w_0^2/4D$ is the diffusion time, and $D$ is the diffusion coefficient. By fitting Eqs. S2-S3 to the experimental autocorrelation function, the diffusion time and amplitudes of the species are obtained. Through $D$, the hydrodynamic diameter $D_H$ can be obtained through the Stokes-Einstein relation.

Figure S1 shows a comparison between sizes obtained with DLS and FCCS. FCCS size data has larger error bars due to the fact that measurements are quasi-single particle, requiring long measurements or multiple measurements to obtain statistical results identical to DLS. Conversely, DLS measurements are more susceptible for bias towards larger averages when samples are polydisperse. The very good agreement between FCCS and DLS results indicates that LPNP samples have low polydispersity[1].

The relation between the amplitude of the autocorrelation function of signal *a* and the number of species contributing to it is given by:

$$A_a = \sum_i \eta_{i,a}{}^2 \cdot N_i \Big/ \left( \sum_i \eta_{i,a} \cdot N_i \right)^2 \tag{S4a}$$

where $N_i$ represents the average number of species *i* in the confocal volume of signal *a*, and $\eta_{i,a}$ represents the brightness of specie $i$, also on the fluorescent signal *a*. Conversely, the cross-correlation amplitude between signals *a* and *b*, $A_X$ is given by:

$$A_X = \sum_i \eta_{i,a} \cdot \eta_{i,b} \cdot N_i \Big/ \left[ \left( \sum_i \eta_{i,a} \cdot N_i \right) \left( \sum_i \eta_{i,b} \cdot N_i \right) \right] \tag{S4b}$$

$A_X$ therefore provides quantitative information regarding the association between the species producing signals *a* and *b*.

### S2. Implementing Fluorescence Cross-Correlation Spectroscopy to quantify formation of DNA-carrying hybrid lipid-polymer nanoparticles

In the present work, polylysine-DNA polyplexes (PPs) are combined with cationic liposomes (Ls) to form hybrid lipid-polymer-DNA nanoparticles (LPNPs). For the simplified case of: (i) one polyplex associating with one cationic liposome (1:1 stoichiometry) to form one LPNP; (ii) no cross-talk between the two channels; and (iii) no changes in the brightness of the species following the association, the equations relating the amplitudes of the correlation curves with the number of species in the confocal volume become straightforward (Eqs 1a-c in the main text)

$$A_G = \left(N_{\mathrm{PP}f} + N_{\mathrm{LPNP}}\right)^{-1} \tag{1a}$$

$$A_R = \left(N_{\mathrm{L}f} + N_{\mathrm{LPNP}}\right)^{-1} \tag{1b}$$

$$A_X = \frac{N_{\mathrm{LPNP}}}{\left(N_{\mathrm{PP}f} + N_{\mathrm{LPNP}}\right) \cdot \left(N_{\mathrm{L}f} + N_{\mathrm{LPNP}}\right)} \tag{1c}$$

Here $A_G$ and $A_R$ are the amplitudes of the green and red auto-correlation, respectively, and $A_X$ is the amplitude of the cross-correlation. $N_{\mathrm{PP}f}$, $N_{\mathrm{L}f}$, and $N_{\mathrm{LPNP}}$, are the average number of free polyplexes, free liposomes and LPNPs in the confocal volumes, respectively.

For the more general case in which one polyplex associates with *n* liposomes to form a $L_n$PNP nanoparticle containing *n* liposomes, and, as before, assuming no cross-talk between the two channels and no changes in the brightness of the species following the association, the equations relating the amplitudes of the correlation curves with the number of species in the confocal volume become:

$$A_G = \frac{1}{\left(N_{\mathrm{PP}f} + N_{\mathrm{LPNP}}\right)} \tag{S5a}$$

$$A_R = \frac{\left(N_{\mathrm{L}f} + n^2 \cdot N_{\mathrm{LPNP}}\right)}{\left(N_{\mathrm{L}f} + n \cdot N_{\mathrm{LPNP}}\right)^2} \tag{S5b}$$

$$A_X = \frac{n \cdot N_{\mathrm{LPNP}}}{\left(N_{\mathrm{PP}f} + N_{\mathrm{LPNP}}\right) \cdot \left(N_{\mathrm{L}f} + n \cdot N_{\mathrm{LPNP}}\right)} \tag{S5c}$$

The fraction of coated $L_n$PNPs is now given by:

$$f_{LPNP} = \frac{{A_X}^2 \cdot (n-1) + A_X \cdot A_G}{n \cdot A_R \cdot A_G} \tag{S6}$$

where $f_{LPNP} = N_{\mathrm{L}_n\mathrm{PNP}} / (N_{\mathrm{PP}f} + N_{\mathrm{L}_n\mathrm{PNP}})$. Note that for *n*=1, Eq. S6 reduces to Eq. 2a. Conversely, the fraction of liposomes that associate with polyplexes continues to be given by the $A_X/A_G$ ratio.

$$\frac{n \cdot N_{\mathrm{L}_n\mathrm{PNP}}}{N_{\mathrm{L}f} + n \cdot N_{\mathrm{L}_n\mathrm{PNP}}} = \frac{A_X}{A_G} \tag{S7}$$

Because both $f_{LPNP}$ and *n* are unknowns, Eq. S6 cannot be used to determine the fraction of coated $L_n$PNPs without knowledge of *n*. However, it can still be used to estimate expected $A_X/A_R$ ratios for different scenarios (i.e. for different fractions of coated $L_n$PNPs and different *n*) and, by comparison with the experimentally observed $A_X/A_R$ ratios, determine which scenarios are more likely.

Fig S2 shows such calculated $A_X/A_R$ ratios using Eqs. S5 and S6 for starting compositions similar to $\rho_{\mathrm{L:DNA}}$=0.75 and 2 (for $\rho_{\mathrm{P:DNA}}$=1.5).

The first important observation is that when the liposome to polyplex ratio ($\rho_N$) is close to 1 (Fig S2a), the deviations between the $A_X/A_R$ ratios and fraction of coated $L_n$PNPs are relatively small (e.g. the largest deviation for $A_X/A_R$=0.8 results in $f_{LPNP}$=0.74 when *n*=1.3), but become greater when $\rho_N$ continues increasing (Fig S2b). This is consistent with the observation that the measured $A_X/A_R$ ratios increase to values greater than 1 as the $\rho_{\mathrm{L:DNA}}$ ratio becomes greater (Fig 2i-j).

The second important observation is that for both compositions illustrated in Fig S2, the experimentally observed $A_X/A_R$ ratios are coincident with estimated $A_X/A_R$ ratios corresponding to $f_{LPNP}$ values of 0.96 or greater, depending on *n*, and *n* values between 1 and 2.3. By also plotting the experimentally observed $A_X/A_G$ ratios, these ranges can be further narrowed down to $f_{LPNP}$≈1 (full conversion), and *n* values of 1 and 1.7. The convergence of both $A_X/A_R$ and $A_X/A_G$ ratios to a scenario of full conversion strongly indicate that the conversion of polyplexes to LPNPs is practically full when the starting polyplexes are negatively charged ($\rho_{\mathrm{P:DNA}}$ = 1 and 1.5), and that the number of liposomes per LPNP is close to 1 in most cases, although it can reach values of 1.7 when $\rho_{\mathrm{L:DNA}}$ becomes high. Lastly, in this calculation, the used $\rho_N$ is estimated by the $A_G/A_R$ ratio (Figure 2k), slightly adjusted to accommodate the fact that *n* can be slightly greater than 1 (e.g. *n*≈1.7 for $\rho_{\mathrm{L:DNA}}$=2), which reveals a good self-consistency between all approximations.

The scenario of full polyplex to LPNP conversion ($f_{LPNP}$≈1) could also be deduced from the fact that deviations between $A_X/A_R$ and $f_{LPNP}$ are small for low $\rho_N$, which then, since $A_X/A_R$=1 for $\rho_{L:DNA}$=0.75 (Figure 2i) indicates full conversion. Hence, further addition of liposomes should not decrease $f_{LPNP}$, which should remain close to complete, indicating that the increase in $A_X/A_R$ most likely reflects a small increase in $n$. If $f_{LPNP}$ is fixed at 1, Eq. S6 can be easily solved, resulting on an average $n$≈1.7 for $\rho_{L:DNA}$=2. This shows that even for the largest $\rho_{L:DNA}$ $n$ is still relatively close to one, which suggests that in general, most polyplexes associate with one liposome, but for greater $\rho_{L:DNA}$, some percentage of polyplexes may associate with more than one liposome.

For $\rho_{P:DNA}$=1 polyplexes (Figure 2j) $A_X/A_R$ increases from ca. 1.25 to 1.4 for $\rho_{L:DNA}$=3 and $\rho_{L:DNA}$=5. This indicates a full conversion of the polyplexes to LPNPs, with an average number of layers $n$ of 1.36 and 1.4, respectively. This is very similar to what is found for the $\rho_{P:DNA}$=1.5 polyplexes, which indicates that despite the larger charge of $\rho_{P:DNA}$=1 polyplexes, a single liposome, which contains ca. 80% of cationic lipid, is sufficient to invert the charge of the polyplexes. The large number of needed liposomes to stabilize the system ($\rho_{L:DNA}$>3) is therefore expected to be mostly needed to neutralize free DNA that was not complexed with the polyplexes.

#### S3. Determining the overlap between the green and red excitation volumes using single- and double-labelled liposomes

The measured cross-correlation amplitude ($A_{X.meas.}$) is limited by the amount of overlap between the green and red excitation volumes. We define the overlap volume correction factor ($V_{crct}$) as the factor that corrects the value of $A_{X.meas.}$ to the value of cross-correlation amplitude ($A_X$) that would be expected if the overlap between the two excitation volumes was perfect, according to Eq. 3,

$$A_X = A_{X.meas.}/V_{crct} \tag{3}$$

To determine $V_{crct}$ and correct the cross-correlation we performed a series of measurements with a set of samples consisting of a mixture between three PEGylated liposomes. Two of the liposomes were labelled with just one dye, one liposome type (L1) with 0.1 mol% of Atto-488 (green), and the other (L2) with 0.1 mol% Texas-red. The third liposome type (L3) was labelled with both dyes simultaneously. By gradually replacing liposomes L1 and L2, which are always in equal amounts and provide a non-colocalized signal, by liposomes L3, in which the two fluorescent probes are expected to be perfectly colocalized, $V_{crct}$ can be determined. A related calibration approach was described recently by Werner et al[2].

The results presented in Figure S3 show that both the $A_{X.meas.}/A_R$ and $A_{X.meas.}/A_G$ ratios increase as the single-labelled liposomes are gradually replaced by liposomes labelled with the two dyes. The maximum cross-correlation percentages detected in the sample with only dual-labeled liposomes were below 80%, due to the non-perfect overlap between the green and red confocal volumes. On the other hand, when the fraction of double-labelled liposomes is zero, there is still some amount of cross-correlation measured, due to the existence of some crosstalk between the dyes. (Note: as an exception to the remaining figures, where cross-correlation amplitudes are always corrected for crosstalk according to the procedure described in Bacia *et al*[3], the data in Figure S3 is not corrected for crosstalk). The expected amplitudes for the auto- and cross-correlation functions in this setting are as follows:

$$A_{\mathrm{G}} = \frac{{\eta_{L1,G}}^2 \cdot N_{L1} + {\eta_{L3,G}}^2 \cdot N_{L3}}{\left(\eta_{L1,G} \cdot N_{L1} + \eta_{L3,G} \cdot N_{L3}\right)^2} \tag{S8a}$$

$$A_{\mathrm{R}} = \frac{{\kappa_{Gr}}^2 \cdot {\eta_{L1,G}}^2 \cdot N_{L1} + {\eta_{L2,R}}^2 \cdot N_{L2} + \left(\kappa_{Gr} \cdot \eta_{L3,G} + \eta_{L3,R}\right)^2 \cdot N_{L3}}{\left(\kappa_{Gr} \cdot \eta_{L1,G} \cdot N_{L1} + \eta_{L2,R} \cdot N_{L2} + \left(\kappa_{Gr} \cdot \eta_{L3,G} + \eta_{L3,R}\right) \cdot N_{L3}\right)^2} \tag{S8b}$$

$$A_{X.meas.} = \frac{V_{crct.} \cdot \left({\eta_{L1,G}}^2 \cdot \kappa_{Gr} \cdot N_{L1} + \eta_{L3,G} \cdot \left(\kappa_{Gr} \cdot \eta_{L3,G} + \eta_{L3,R}\right) \cdot N_{L3}\right)}{\left(\eta_{L1,G} \cdot N_{L1} + \eta_{L3,G} \cdot N_{L3}\right) \cdot \left(\kappa_{Gr} \cdot \eta_{L1,G} \cdot N_{L1} + \eta_{L2,R} \cdot N_{L2} + \left(\kappa_{Gr} \cdot \eta_{L3,G} + \eta_{L3,R}\right) \cdot N_{L3}\right)} \tag{S8c}$$

where $\kappa_{Gr}$ is a factor describing the fraction of signal from the green liposomes (L1) that is detected in the red detector due to crosstalk. Both $A_{\mathrm{X}}/A_{\mathrm{R}}$ and $A_{\mathrm{X}}/A_{\mathrm{G}}$ values are fitted simultaneously with Eq. S8, allowing a robust determination of $V_{crct} = 0.746$.

Besides providing the correction factor for the volume overlap, this experiment also validates the suitability of FCCS to determine the colocalization of soft nanoparticles of ca. 100 nm.

**S4. LPNP assembly behaviour and ternary map**

Figure 4 summarises the results and observations of this work in the form of a ternary map of assembly in excess water (given the out-of-equilibrium nature of the system, the map is not strictly a phase diagram).

The first important observation is the lack of formation of LPNPs when starting with positive ($\rho_{\mathrm{P:DNA}} = 3$) polyplexes (region C on the map). It is unclear whether replacing polylysine with a weaker polycation (e.g., polyethylenimine) could result in some displacement of the polymer to allow the deposition of cationic liposomes and enable LPNP formation.

The second important observation is the need to have at least one liposome per polyplex ($\rho_{\mathrm{N}} \geq 1$) to form stable LPNPs. This ensures that every polyplex is enveloped by a liposome, having their negative charge reversed and inhibiting aggregation. This requirement divides region B into two subregions: B1, where aggregation is observed, and B2, where LPNPs stable for at least one week are formed. For $\rho_{\mathrm{P:DNA}} = 1.5$, the $\rho_{\mathrm{N}} = 1$ boundary was estimated at $\rho_{\mathrm{L:DNA}} = 0.63$ (Figure 2k). For $\rho_{\mathrm{P:DNA}} = 1$, the boundary is difficult to estimate because of the excess DNA, but it lies between $\rho_{\mathrm{L:DNA}} = 3$ and $5$. As an approximation the B1-B2 boundary is thus drawn as a straight line starting in the isoelectric point ($\rho_{\mathrm{P:DNA}} = 1.6$, $\rho_{\mathrm{L:DNA}} = 0$) and intersecting the point $\rho_{\mathrm{P:DNA}} = 1.5$, $\rho_{\mathrm{L:DNA}} = 0.63$. Besides the approximation, since both the number of polyplexes and liposomes depend both on composition and preparation method, the boundary line is specific to the present work, but is still insightful, and can be used as a point of departure when working with other LPNP systems.

In B2, where stable LPNPs are formed, the conversion of polyplexes to LPNPs is practically full, indicating that neither the polycation nor DNA are displaced by the liposomes. However, the LPNPs still coexist with other (non-competing) species. Along the $\rho_{\mathrm{P:DNA}} = 1.5$ line LPNPs coexist with excess liposomes, whereas for $\rho_{\mathrm{P:DNA}} = 1$, they almost certainly coexist with lipoplexes resulting from the complexation of excess DNA. Hence, when investigating biologic activity of LPNP systems, especially if away from $\rho_{\mathrm{iso}}$, the possible coexistence with lipoplexes should be taken into consideration in the elucidation of transfection and therapeutic potential of the formulations.

We have also observed that $\rho_{\mathrm{P:DNA}} = 1$ systems form ca. 3.5 times more polyplexes than $\rho_{\mathrm{P:DNA}} = 1.5$ do. This suggests that the polymer-DNA cores contain a smaller number of DNA plasmids in the former than in the latter, which should be taken into consideration since the number of encapsulated plasmids can be therapeutically relevant[4].


**Figures and Tables**

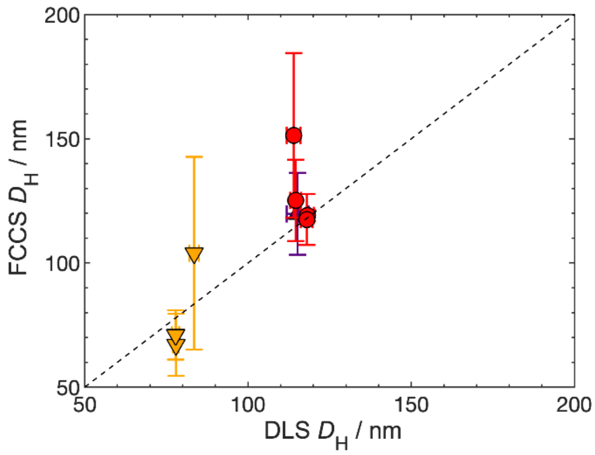


**Figure S1**. Comparison between the LPNPs' hydrodynamic diameter ($D_H$) obtained with DLS and FCCS. The differences between sizes determined with DLS and FCCS are relatively small, indicating relatively low sample polydispersity[1].

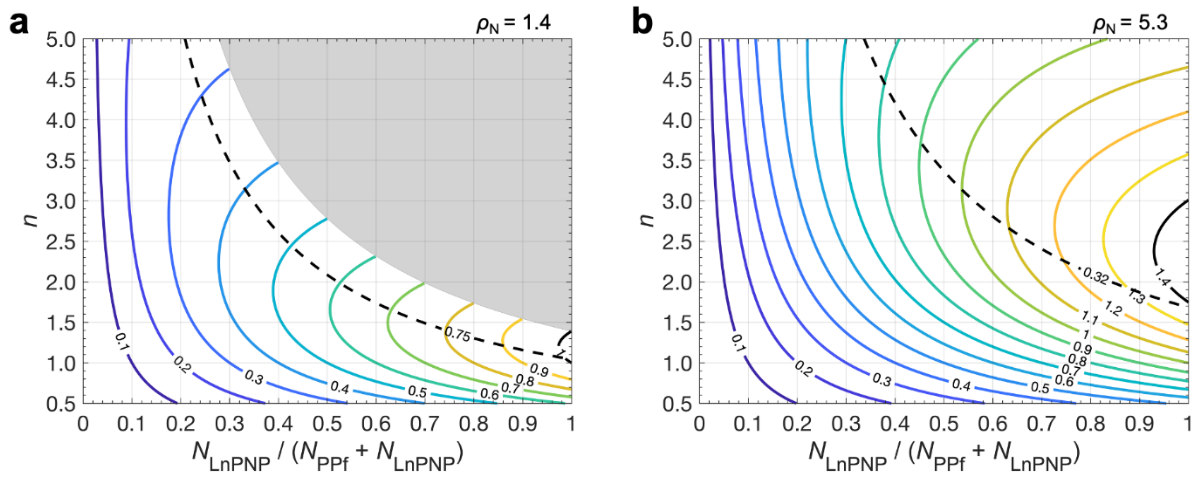


**Figure S2**. Contour map for the estimated $A_X/A_R$ ratios for different values of conversion rates ($f_{LPNP} = N_{\mathrm{LPNP}}/(N_{\mathrm{PP}f} + N_{\mathrm{LPNP}})$), number of liposomes per L$_n$PNP ($n$), and liposome to polyplex ratio ($\rho_N$). The different colored lines indicate compositions with equal $A_X/A_R$ ratios. For $n$=1, the $A_X/A_R$ ratio is equal to $f_{LPNP}$, as dictated by Eq. 2a. The shaded region encompasses inaccessible LPNP compositions, for which the fraction of used liposomes would be above 1. For $\rho_N$=1.4 (**a**), which is a composition close to $\rho_{\mathrm{L:DNA}}$=0.75, $\rho_{\mathrm{P:DNA}}$=1.5, the experimentally measured $A_X/A_R$ ratio is 1 (signaled by the black straight curve). Under these conditions, the expected system composition indicates a conversion ratio between 1 (for $n$=1) and 0.97 (for $n$=1.2). For $\rho_N$=5.3 (**b**), which is a composition close to $\rho_{\mathrm{L:DNA}}$=2, $\rho_{\mathrm{P:DNA}}$=1.5, the experimentally measured $A_X/A_R$ ratio is 1.4 (signaled by the black curve). Under these conditions, the expected system composition indicates a conversion ratio between 1 (for $n$=1.7) and 0.96 (for $n$=2.3). Both (a) and (b) show parabola type curves for $A_X/A_R$, indicating that more than one solution for a single conversion fraction could exist, but the lower bounds are more consistent with the experimentally measured $A_X/A_G$ ratios (signaled by the black dashed curves).

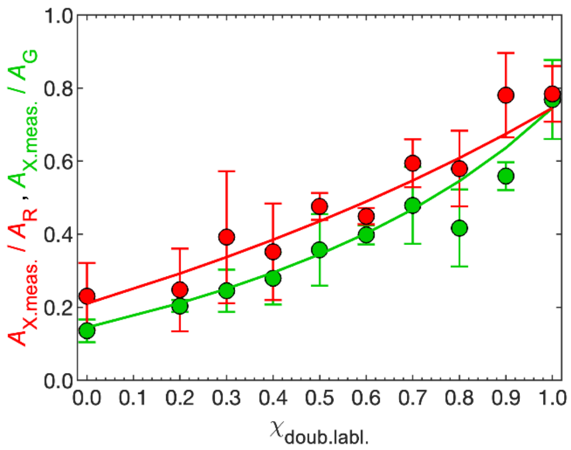


**Figure S3.** Determination of the confocal overlap volume $V_{crct}$. By employing a mixture of liposomes labelled with either Atto-488 or Texas Red, and liposomes labelled with both dyes, the fraction of colocalization in solution can be controlled and compared with the measured cross-correlation. As the fraction of dual-labelled liposomes increases, $A_{X.meas.}/A_G$ (green symbols) and $A_{X.meas.}/A_R$ (red symbols) also increase, as expected. The estimated $V_{crct}$, obtained by fitting both curves simultaneously, is 74.6%. The measurements also confirm the suitability of FCCS to measure colocalization in soft self-assembled nanostructures of ca. 100 nm.

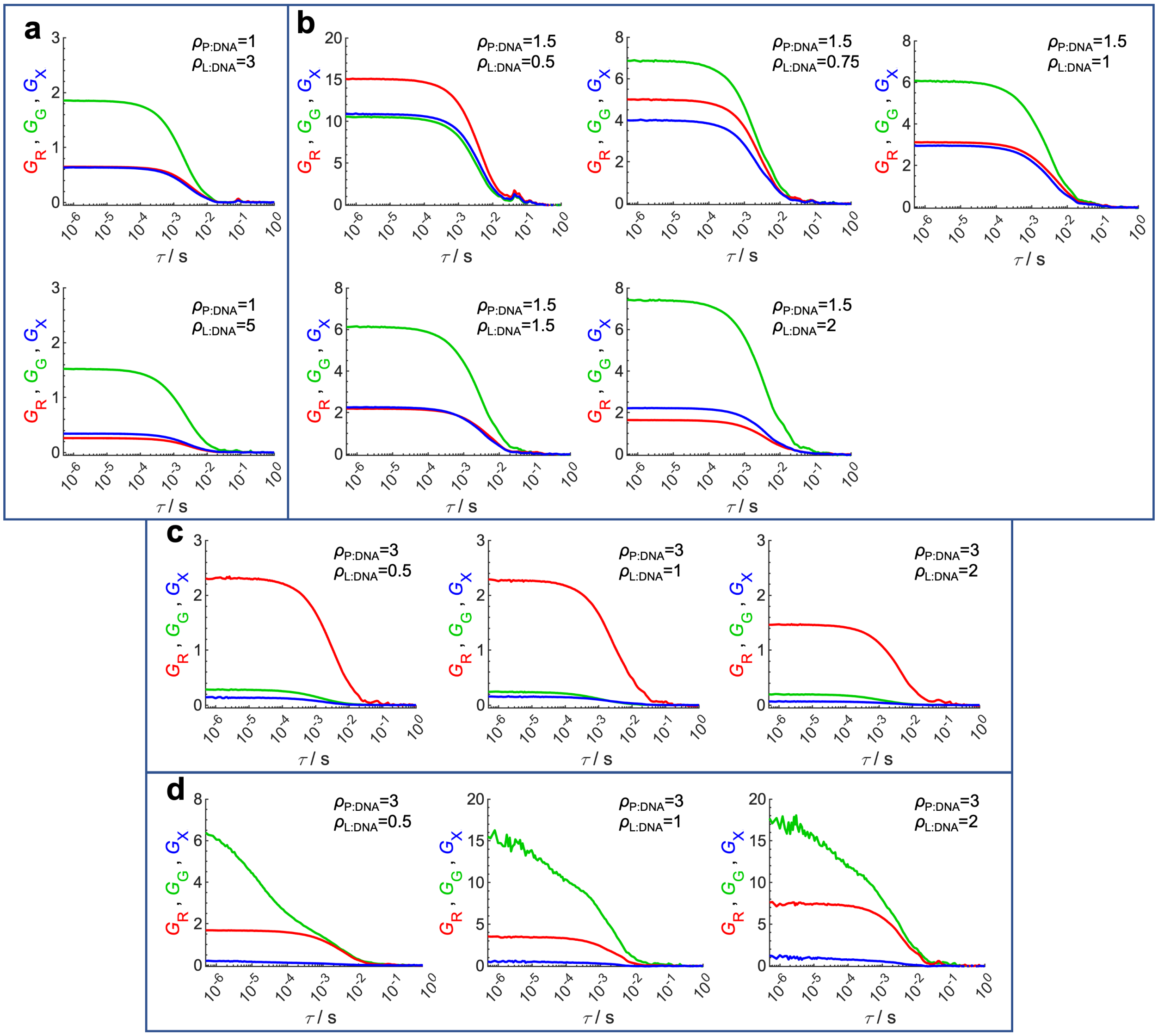

**Figure S4**. Representative auto- and cross-correlation curves measured for each of the LPNP formulations using Texas Red-labelled cationic liposomes with 10% PEG. (**a-c**) LPNPs using polyplex cores with Atto 488-labeled polylysine ($\rho_{\mathrm{P:DNA}} = 1,\ 1.5$ and $3$). (**d**) LPNP formulations prepared employing a cationic polyplex core ($\rho_{\mathrm{P:DNA}} = 3$), where the DNA is labelled with YOYO-1. Green auto-correlation curves are shown in green, red auto-correlation curves are shown in red, and cross-correlation curves are shown in blue. The existence of labelled free polylysine in excess in the $\rho_{\mathrm{P:DNA}} = 3$ system lowers dramatically the amplitude of the green autocorrelation to values where it becomes inaccurate. This hinders the determination of the association between cationic polyplexes and cationic liposomes (**c**). Hence, in this case, DNA was also labelled with YOYO-1 (**d**). The very low values of cross-correlation amplitude compared to both the autocorrelation amplitudes shows that cationic polyplexes and cationic liposomes do not associate. Note that if the amplitude of the cross-correlation was large, such result would not be able to confirm association between polyplexes and liposomes, since DNA could also be displaced to the liposomes, but the absence of cross-correlation is sufficient to rule out association.

**Table S1.** Characterization by DLS and ζ Potential of polyplexes prepared with different charge ratios. Data are Means ± SD (N=3).

| $\rho_{P:DNA}$ | $D_H$ (Z Average) / nm | PDI | ζ Potential /mV |
|---|---|---|---|
| **1** | 120.7 ± 2.2 | 0.37 ± 0.02 | -69 ± 3.5 |
| **1.5** | 133.8 ± 3.4 | 0.29 ± 0.01 | -36 ± 2.7 |
| **3** | 79 ± 3.1 | 0.32 ± 0.03 | +38.9 ± 1.8 |

**Table S2**. Characterization by DLS and ζ Potential of cationic liposomes prepared with different PEGylation degrees. Data are Means ± SD (N=3).

| PEG mol% | $D_H$ (Z Average) / nm | PDI | ζ Potential /mV |
|---|---|---|---|
| **10** | 99.2 ± 2.4 | 0.46 ± 0.034 | +128.6 ± 2 |
| **5** | 89.1 ± 5.8 | 0.44 ± 0.051 | +120.6 ± 5.5 |
| **0** | 91.9 ± 2.9 | 0.41 ± 0.044 | +62.3 ± 1.5 |

**Table S3**. Characterization by DLS and ζ Potential of LPNP formulations prepared employing different polyplexes ($\rho_{P:DNA}$=1, 1.5 and 3) combined with 10 mol% cationic liposomes. Data are Means ± SD (N=3).

| $\rho_{P:DNA}$ | $\rho_{L:DNA}$ | $D_H$ (Z Average) / nm | PDI | ζ Potential /mV |
|---|---|---|---|---|
| 1 | 0.5 | 12184 ± 3269 | 17.71 ± 7.14 | +17.8 ± 0.7 |
| 1 | 1 | 782.4 ± 336.5 | 1.487 ± 0.40 | +35.9 ± 2.5 |
| 1 | 2 | 686.8 ± 19.2 | 2.46 ± 0.02 | +48 ± 2.9 |
| 1 | 3 | 341.1 ± 143.8 | 1.003 ± 0.687 | +62.4 ± 3 |
| 1 | 5 | 115.2 ± 3.3 | 0.377 ± 0.01 | +65 ± 3.5 |
| 1.5 | 0.5 | 331.6 ± 372.77 | 0.8425 ± 1.327 | +21.9 ± 5.7 |
| 1.5 | 0.75 | 118.2 ± 2.0 | 0.229 ± 0.02 | +30.0 ± 8.6 |
| 1.5 | 1 | 114.6 ± 1.7 | 0.209 ± 0.037 | +40.6 ± 1.8 |
| 1.5 | 1.5 | 114 ± 2.1 | 0.228 ± 0.009 | +49.4 ± 0.4 |
| 1.5 | 2 | 118 ± 1.8 | 0.3 ± 0.043 | +57 ± 0.8 |
| 3 | 0.5 | 77.9 ± 0.8 | 0.26 ± 0.03 | +50.2 ± 0.7 |
| 3 | 1 | 77.8 ± 1.1 | 0.27 ± 0.03 | +51.6 ± 2.7 |
| 3 | 2 | 83.5 ± 1.5 | 0.33 ± 0.03 | +53.3 ± 0.1 |

**Table S4**. Size characterization by FCCS of LPNP formulations prepared employing different polyplexes ($\rho_{P:DNA}$=1, 1.5 and 3) combined with cationic liposomes (10% PEG). Data are Means ± SD (N=3).

| $\rho_{P:DNA}$ | $\rho_{L:DNA}$ | Diffusion coefficient / µm$^2$ s$^{-1}$ | $D_H$ (FCCS) / nm |
|---|---|---|---|
| 1 | 5 | 4.15 ± 0.55 | 119.79 ± 16.53 |
| 1.5 | 0.5 | 3.37 ± 0.66 | 149.31 ± 26.64 |
| 1.5 | 0.75 | 4.98 ± 1.70 | 119.13 ± 2.34 |
| 1.5 | 1 | 3.97 ± 0.50 | 125.19 ± 16.33 |
| 1.5 | 1.5 | 3.34 ± 0.66 | 151.35 ± 33.13 |
| 1.5 | 2 | 4.20 ± 0.35 | 117.49 ± 10.21 |
| 3 | 0.5 | 7.51 ± 1.41 | 67.01 ± 12.53 |
| 3 | 1 | 7.00 ± 0.92 | 71.06 ± 9.93 |
| 3 | 2 | 5.16 ± 1.75 | 103.94 ± 38.83 |

**Table S5**. Summary of FCCS data for LPNPs. Results shown are obtained after fitting the auto- and cross-correlation curves with Eqs. 1 and 2. Data are Means ± SD (N≥3).

| $\rho_{P:DNA}$ | $\rho_{L:DNA}$ | $A_G$ | $A_R$ | $A_X$ | $A_G^{-1}$ [a] | $A_R^{-1}$ [a] | $A_X$ / $A_R$ [b] | $A_X$ / $A_G$ [b] |
|---|---|---|---|---|---|---|---|---|
| | | | | **0 mol% PEG** | | | | |
| 1.5 | 0.5 | 8.97 ± 0.62 | 10.01 ± 0.50 | 10.55 ± 0.29 | 0.112 ± 0.008 | 0.100 ± 0.005 | 1.06 ± 0.04 | 1.18 ± 0.09 |
| 1.5 | 0.75 | 5.9 ± 2.3 | 4.78 ± 0.88 | 5.3 ± 1.7 | 0.191 ± 0.069 | 0.216 ± 0.044 | 1.08 ± 0.18 | 0.907 ± 0.086 |
| 1.5 | 1 | 7.9 ± 2.8 | 4.38 ± 0.81 | 5.2 ± 1.3 | 0.150 ± 0.083 | 0.235 ± 0.041 | 1.20 ± 0.28 | 0.70 ± 0.11 |
| 1.5 | 1.5 | 5.9 ± 1.3 | 2.74 ± 0.72 | 3.10 ± 0.66 | 0.177 ± 0.050 | 0.383 ± 0.084 | 1.15 ± 0.18 | 0.53 ± 0.10 |
| 1.5 | 2 | 7.0 ± 2.0 | 1.93 ± 0.40 | 2.67 ± 0.55 | 0.156 ± 0.058 | 0.54 ± 0.12 | 1.43 ± 0.39 | 0.40 ± 0.12 |
| | | | | **5 mol% PEG** | | | | |
| 1.5 | 0.5 | 10.43 ± 0.56 | 13.2 ± 1.0 | 13.28 ± 0.77 | 0.096 ± 0.005 | 0.076 ± 0.006 | 1.01 ± 0.03 | 1.27 ± 0.01 |
| 1.5 | 0.75 | 6.4 ± 2.9 | 5.0 ± 1.7 | 5.1 ± 1.7 | 0.185 ± 0.079 | 0.220 ± 0.072 | 1.02 ± 0.15 | 0.83 ± 0.12 |
| 1.5 | 1 | 8.0 ± 3.1 | 4.48 ± 0.84 | 4.82 ± 0.46 | 0.137 ± 0.037 | 0.231 ± 0.048 | 1.10 ± 0.17 | 0.65 ± 0.16 |
| 1.5 | 1.5 | 6.0 ± 1.8 | 2.9 ± 1.3 | 3.2 ± 1.1 | 0.186 ± 0.083 | 0.42 ± 0.24 | 1.13 ± 0.14 | 0.53 ± 0.11 |
| 1.5 | 2 | 6.8 ± 3.1 | 1.99 ± 0.75 | 2.44 ± 0.81 | 0.19 ± 0.13 | 0.57 ± 0.22 | 1.25 ± 0.27 | 0.40 ± 0.13 |
| | | | | **10 mol% PEG** | | | | |
| 1 | 3 | 1.81 ± 0.09 | 0.52 ± 0.13 | 0.652 ± 0.086 | 0.553 ± 0.027 | 1.98 ± 0.45 | 1.27 ± 0.16 | 0.360 ± 0.038 |
| 1 | 5 | 1.68 ± 0.14 | 0.32 ± 0.09 | 0.419 ± 0.035 | 0.598 ± 0.053 | 3.26 ± 0.82 | 1.35 ± 0.26 | 0.250 ± 0.013 |
| 1.5 | 0.5 | 10.4 ± 1.5 | 14.9 ± 1.8 | 14.3 ± 1.6 | 0.097 ± 0.015 | 0.068 ± 0.009 | 0.962 ± 0.044 | 1.37 ± 0.05 |
| 1.5 | 0.75 | 7.3 ± 4.3 | 5.2 ± 1.5 | 5.2 ± 2.1 | 0.171 ± 0.078 | 0.207 ± 0.055 | 0.98 ± 0.15 | 0.75 ± 0.10 |
| 1.5 | 1 | 7.1 ± 3.4 | 3.46 ± 0.58 | 3.7 ± 1.1 | 0.19 ± 0.14 | 0.295 ± 0.043 | 1.09 ± 0.32 | 0.58 ± 0.12 |
| 1.5 | 1.5 | 7.9 ± 3.7 | 3.5 ± 3.0 | 3.6 ± 2.2 | 0.147 ± 0.062 | 0.40 ± 0.16 | 1.19 ± 0.30 | 0.447 ± 0.077 |
| 1.5 | 2 | 6.5 ± 1.4 | 1.46 ± 0.19 | 2.02 ± 0.62 | 0.161 ± 0.040 | 0.696 ± 0.093 | 1.42 ± 0.48 | 0.314 ± 0.080 |
| 3 | 0.5 | 7.3 ± 5.4 | 5.6 ± 2.2 | 0.43 ± 0.19 | 0.31 ± 0.37 | 0.199 ± 0.073 | 0.076 ± 0.019 | 0.088 ± 0.062 |
| 3 | 1 | 7.8 ± 5.4 | 3.00 ± 0.58 | 0.24 ± 0.13 | 0.28 ± 0.33 | 0.342 ± 0.061 | 0.079 ± 0.044 | 0.039 ± 0.018 |
| 3 | 2 | 2.04 ± 0.14 | 1.73 ± 0.51 | 0.105 ± 0.019 | 0.492 ± 0.034 | 0.62 ± 0.19 | 0.062 ± 0.009 | 0.052 ± 0.012 |

[a]Note that for the simplified model of one liposome complexing with one polyplex (1:1 stoichiometry), $A_G^{-1}$ and $A_R^{-1}$ correspond to the total number of polyplexes ($N_{PPf} + N_{LPP}$) and liposomes ($N_{Lf} + N_{LPP}$) per confocal volume in the sample, respectively. [b]Note that within the same 1:1 stoichiometry approximation $A_X/A_R$ and $A_X/A_G$ indicate the fraction of polyplexes converted to LPNPs ($f_{LPNP}$) and fraction of liposomes used in LPNPs, respectively.
